# Detection of latent brain states from baseline neural activity in the amygdala

**DOI:** 10.1101/2024.06.14.598974

**Authors:** Alexa Aucoin, Kevin K. Lin, Katalin M. Gothard

## Abstract

The amygdala responds to a large variety of socially and emotionally salient environmental and interoceptive stimuli. The context in which these stimuli occur determines their social and emotional significance. In canonical neurophysiological studies, the fast-paced succession of stimuli and events induce phasic changes in neural activity. During inter-trial intervals neural activity is expected to return to a stable and relatively featureless baseline. Context, such as the presence of a social partner, or the similarity of trials in a blocked design, induces brain states that can transcend the fast-paced succession of stimuli and can be recovered from the baseline firing rate of neurons. Indeed, the baseline firing rates of neurons in the amygdala change between blocks of trials of gentle grooming touch, delivered by a trusted social partner, and non-social airflow stimuli, delivered by a computer-controlled air valve. In this experimental paradigm, the presence of the groomer alone was sufficient to induce small but significant changes in baseline firing rates. Here, we examine local field potentials (LFP) recorded during these baseline periods to determine whether context was encoded by network dynamics that emerge in the local field potentials from the activity of large ensembles of neurons. We found that machine learning techniques can reliably decode social vs. non-social context from spectrograms of baseline local field potentials. Notably, decoding accuracy improved significantly with access to broad-band information. No significant differences were detected between the nuclei of the amygdala that receive direct or indirect inputs from areas of the prefrontal cortex known to coordinate flexible, context-dependent behaviors. The lack of nuclear specificity suggests that context-related synaptic inputs arise from a shared source, possibly interoceptive inputs that signal the sympathetic- vs. parasympathetic-dominated states characterizing non-social and social blocks, respectively.

## INTRODUCTION

During natural engagement with the environment, the brain concomitantly processes stimuli that convey specificity to an event and the context in which the event occurs. Stimulus parameters, internal transformations, and the resulting behaviors can be decoded from the activity of ensemble of simultaneously active neurons, and local field potentials (LFPs). Decoding context, however, is challenging because, in most experimental settings, context is unchanging and often conflated with stimulus-evoked neural activity. Context signaling is more likely confined to baseline activity, as demonstrated by neurophysiological studies that independently varied the behaviorally relevant stimuli and the context. For example, when rats learned the probability of a predator interfering with their run toward a coveted reward, baseline activity in the amygdala exhibited correlation with the likelihood of an encounter with the predator. While baseline firing rate varied in proportion to their anticipatory anxiety, the predator-induced firing rate remained unchanged (Amir et al., 2019). In a similar vein, when monkeys learned to associate odors with positive and negative outcomes, the baseline firing rate of neurons in the amygdala and anterior cingulate cortex retained information about the strength of the learned association during the time intervals between trials (Taub et al., 2018). Beyond the amygdala and affective states, the baseline activity of neurons in the neocortex and the basal ganglia can retain information about the outcome of numerous preceding trials that contribute to outcome predictions for upcoming trials (Histed et al., 2019). Likewise, baseline neural activity in the marmoset prefrontal cortex preceding and following a perceived vocalization predicted the likelihood of a reciprocating response (Jovanovic et al., 2022).

It appears, therefore, that neural activity during baseline is fertile ground to explore how the brain might integrate stimuli and events across multiple time scales, how it predicts – rather than reacts to – external events, and how it creates persistent affective states. Indeed, affective states, such as anxiety, persist longer than an emotional reaction to the negatively or positively valanced external stimulus. We have recently demonstrated that grooming, the most common form of social and affective touch in macaques, elicits persistent changes in baseline firing rate in 25-45% of neurons in the amygdala (Martin et al., 2023). The observed changes in baseline were correlated with the animal’s physiological state (low sympathetic and high parasympathetic tone) and with the social context. The presence of the groomer near the monkey, even in the absence of grooming gestures, was sufficient to shift the baseline in the direction in which grooming would shift it. However, this earlier study was focused on the baseline firing rates of individual neurons and left open the possibility that the joint activity of neural populations at the mesoscale level contains comparable information about the brain state of the animal. This is significant because (1) the presence of such information in the LFPs would indicate that contextual information is not just carried by select neurons but reflects a wider state change in the relevant circuitry; (2) from an experimental point of view, LFPs are more robust than single units and less prone to processing artifacts. In this paper, we examine the encoding of contextual information in baseline LFPs, focusing on three questions.

First, we asked whether social context can be decoded from LFP recorded during intertrial intervals (baseline activity) from the amygdala. The large fraction of neurons (25-45%) that showed context-dependent changes in baseline firing rates gives rise to a specific covariance pattern across a population of neurons. Such covariance patterns, or “latent dynamics” have been detected in both single unit activity and LFPs (Gallego-Carracedo et al., 2022) but only during engagement with a stimulus or task parameter. We hypothesize that these latent dynamics are context-dependent, can persist across trials, and can be decoded from baseline activity.

Second, we asked whether there is a detectable difference in data recorded from different nuclei of the amygdala. We hypothesized that context-related activity will be most prominent in the basal and accessory basal nuclei that receive more robust inputs from the prefrontal cortex than the lateral and central nuclei (McDonald 1998, Pitkänen and Amaral, 1991; Barbas 2007; Price and Amaral 1981). As the LFP in each nucleus (subjected to common reference averaging to eliminate volume-conducted components), result from the synaptic currents received and summed across thousands of neurons (Buzsáki et al., 2012; Pesaran et al., 2018), the LFP in the basal and accessory basal nuclei may be driven by inputs from the primate-specific areas of the prefrontal cortex that enable context-dependent flexible emotional behaviors (Passingham and Wise, 2012). Alternatively, the context-related activity may arise from interoceptive inputs, signaling to the brain the parasympathetic-dominated physiological state observed during the grooming blocks (Martin et al, 2023). In this case, we expect comparable decoding accuracy from all nuclei of the amygdala.

Third, we asked whether contextual information is encoded in specific frequency bands, or if it is distributed across multiple frequency components. Localization in frequency domain may be indicative of synchronous activity; such activity may also serve to further coordinate neural activity, by recruiting other populations.

To detect contextual information in LFP, we use modern machine learning (ML) techniques in combination with a simple cross-validation procedure to test whether baseline LFP contains contextual information. The accuracy of the trained classifier can be viewed as measuring the degree to which context can be inferred from LFP.

## RESULTS

### 1. Experimental design

Three adult male macaques received, in alternating blocks of trials, two types of tactile stimuli: (1) a gentle airflow (not a startling air puff) with a pressure of 10 Pa delivered through airflow nozzles brought to the vicinity of the face, but avoiding the eyes and the nostrils, and (2) gentle grooming sweeps delivered to the same areas of the face by a trusted human partner, who wore an instrumented glove that allowed matching the contacts forces of the airflow and the grooming sweeps (**Figure 1**). Linear electrode arrays (V-probes) with 32 recording contacts distributed across a span of 6mm from the tip were lowered into the amygdala. The 6mm-span ensured that we recorded LFP from the full dorso-ventral expanse of the amygdala. On each recording session the V-probes were lowered to different anterior-posterior and medial-lateral coordinates of the amygdala to enable quasi-equal sampling of all component nuclei. The location of the recording electrode was determined through MRI reconstruction (**Figure 1E**). During grooming blocks, the heart rates of the subjects were significantly reduced compared to airflow blocks, indicating a state of low sympathetic arousal (**Figure 2A**). Moreover, heart rate variability was increased during grooming, which is a reliable sign of parasympathetic-dominated physiological state (Berntson et al., 1003) (**Figure 2B**). Respiratory sinus arrhythmia (RSA) taken over 60s = intervals show statistically higher strength during grooming than airflow in two of three subjects. (One-sided t-test: Monkey A, p = 0.0018, n=12 sessions; Monkey S, p < 0.001, n=8 sessions; Monkey C, p > 0.05, n=9 sessions.) Each airflow sequence consists of 11 presentations of the stimulus to pseudo-random locations (10 aimed at the face, one sham). Each presentation of 1s duration is separated by 4s. During a block, the sequence is repeated 10 times for a total of 110 presentations. Each grooming block consists of 20 stimuli, separated by ∼4s and repeated 5 times for a total of 100 presentations. A few minutes elapsed between blocks.

**Figure 1.**
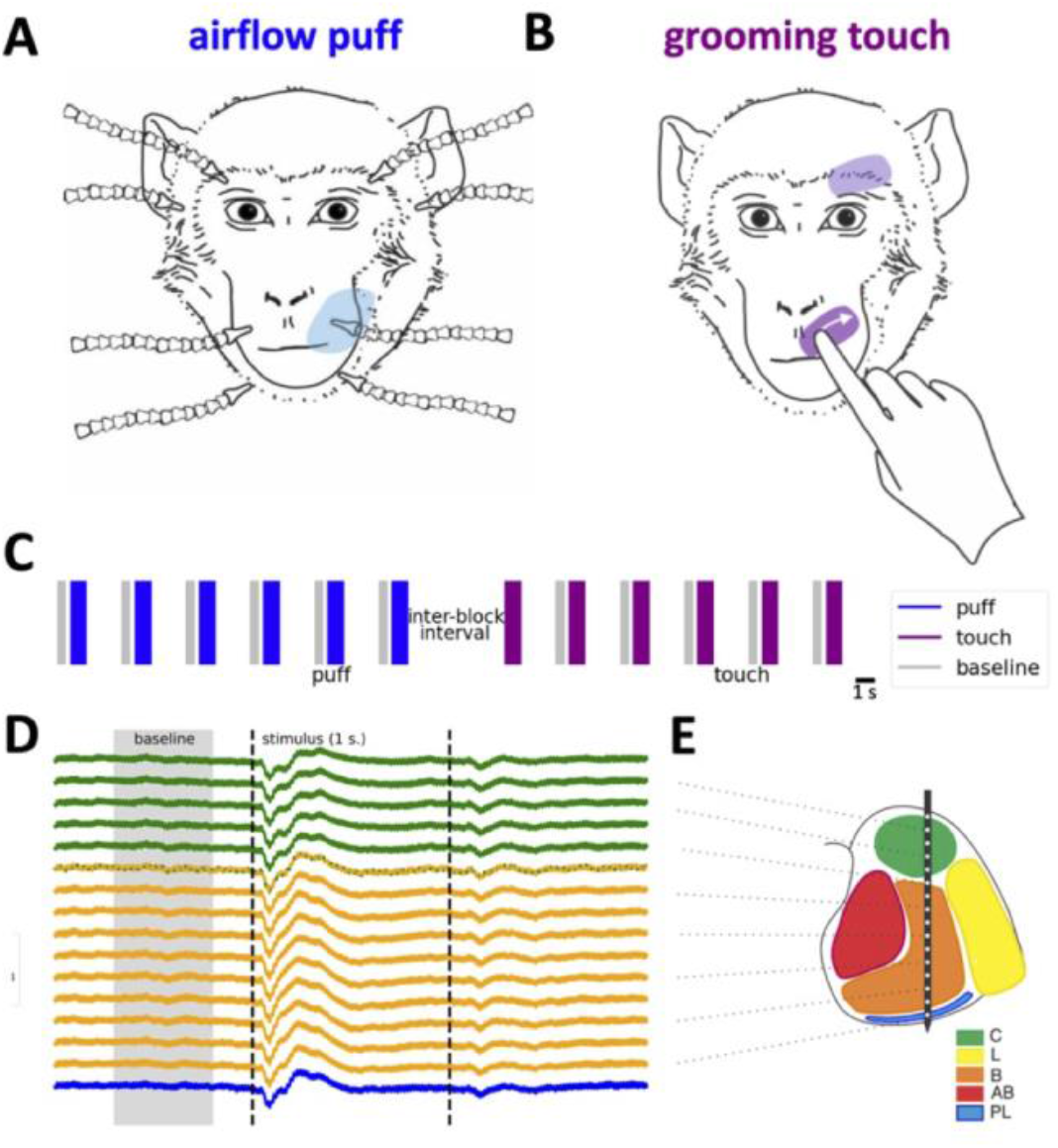
Experimental design. **(A)** Gentle airflow was delivered by custom-built system of air nozzles supplied by computer-controlled pressure valves that produced airflows of 10 Pa for a duration of 1 s. The shaded area on the right upper muzzle indicates the spread of skin mechanoceptors activated by the stimulus. **(B)** Grooming sweeps to the same skin area as in (A) delivered by a trusted human. **(C)** Time course of the last 6 airflow trials in an airflow block followed by the first 6 grooming trials in the subsequent touch block. Blue and maroon vertical lines indicate successive airflow and touch trials, respectively. The width of the line indicates the stimulus duration = 1s. Vertical gray bars indicate the baseline selected between two stimuli of the same type. Note that there is no baseline selection before the first trial of a new block. **(D)** Event-related LFP from a sample recording session. Color code of LFP activity corresponds to the estimated location of V-probe contacts in different nuclei of the amygdala. Lines with alternating colors refer to contacts on the boundary of two nuclei (**E)**. Corresponding recording sites in the amygdala. C=central, green; L= lateral, yellow; B = basal, orange; AB = accessory basal, red; Pl = paralaminar, blue.

**Figure 2.**
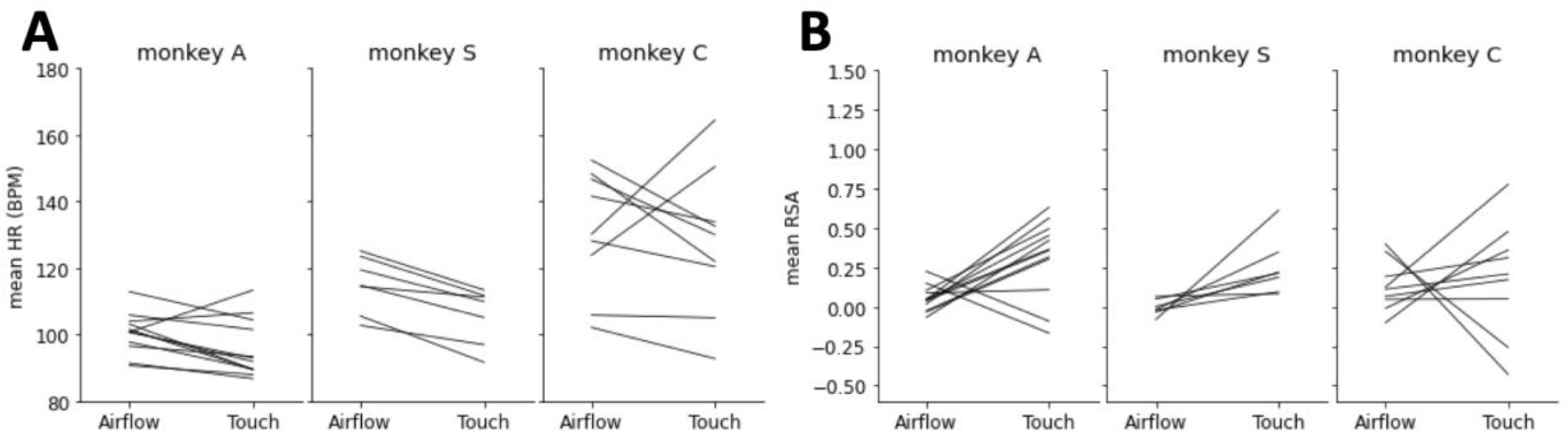
Autonomic state difference in airflow versus grooming blocks. (A) Mean heart rate measurements during airflow and grooming blocks for Monkey A (left), Monkey S (center), and Monkey C. (B) Mean RSA strength during airflow and grooming blocks for Monkey A (left), Monkey S (center), and Monkey C.

### 2. Baseline selection criteria

Baseline LFPs were selected from stable time windows of the interstimulus interval (ISI) between two stimuli of the same type. In other words, we did not consider baseline trials occurring before the first stimulus presentation in each block. The ISI was defined as the period occurring 200ms after stimulus offset and 200ms before stimulus onset. Baseline time windows were selected from ISI for their stable statistical properties. ISI signals in each session were trial averaged and the standard deviation for each timepoint was calculated. Baseline LFP for each trial was chosen by inspecting the trial-averaged ISI and determining a time window with low trial-wise variability. See SI for details.

### 3. Machine Learning for LFP analysis

Recent years have seen a number of major advances in ML in biomedical sciences, including cancer diagnosis (Kourou et. al. 2017), detection and treatment of Alzheimer’s and Parkinson’s diseases (Moradi et. al. 2015, Golshan et. al 2020), and seizure detection (Zhou et. al. 2018). We are particularly motivated by LFP-NET (Golshan et. al 2020), which uses convolutional neural networks to analyze LFP data from human subjects with DBS implants. We implemented our ML-based methodology using two well-known and popular types of classifiers: a convolutional neural network (CNN) based on (Golshan et. al 2020), and a support vector machine (SVM) (Pedregosa et al 2011) (see Materials & Methods). The use of two classifiers allows us to check our findings and compare their performance in a practical setting (see SI for more details on ML architectures and comparisons). Spectral features are often chosen as a reliable feature space for decoding behavior from LFP (Angjelichinoski et al., 2019). To make use of spectral information and at the same time accommodate potential nonstationarity in the data, we use single trial (∼500ms) time-frequency plots, or spectrograms, of LFP as inputs to our classifiers. Deep neural networks have been rarely applied to characterize single trial LFP events (Shilling et al., 2022). Both CNN and SVM leverage statistical methods to nonlinearly transform baseline spectrograms and “learn” spatiotemporal patterns (features) that separate airflow from touch in this new feature space. Given that context-related modulation was seen in a fraction of the single units (Martin et al., 2023), we hypothesize that baseline LFP, which records from a larger population of neurons at the mesoscale, may provide a more reliable feature space for successfully decoding context than single-units.

The workflow, the format of the data, and features of the two classifiers used to decode baseline trials are shown in **Figure 3**. For each recording session and for each nucleus recorded from on that session, we trained one CNN and one SVM. Baseline trials were labelled as “airflow” (for neural activity occurring during the baseline between two airflow trials in an airflow block) or “touch” (for neural activity occurring between two grooming trials in a grooming block).

**Figure 3.**
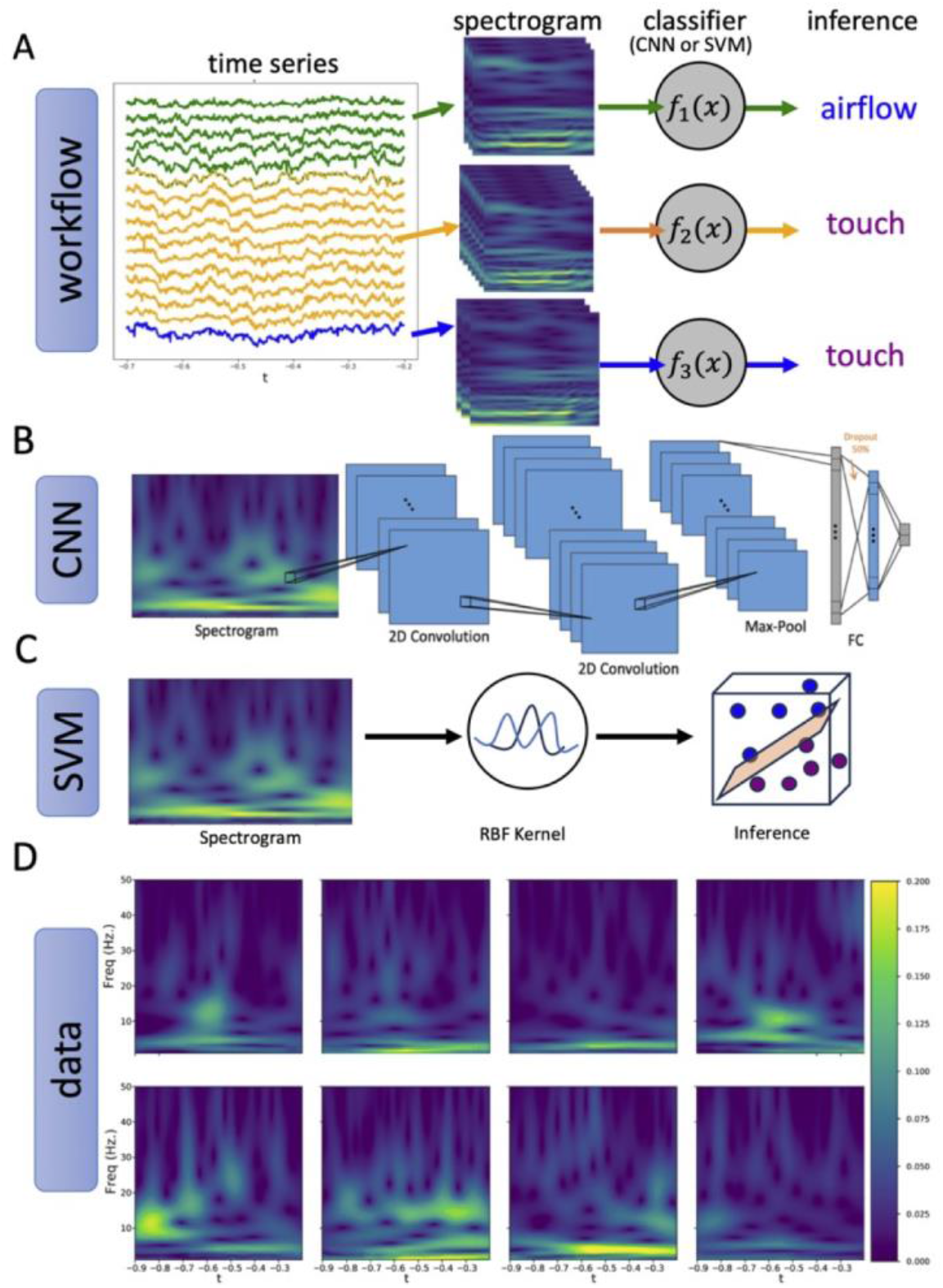
Analysis pipeline. **(A)** LFP trace from a single baseline trial. Signals recorded from the same nucleus (indicated by the same color) are grouped together. Spectrograms are computed using a complex Morlet wavelet transform and labeled as “airflow” or “touch” depending on block type, then used to train a classifier (CNN or SVM) to discriminate between airflow and touch baseline for each nucleus. **(B)** Architecture of the CNN consisting of two 2D convolutional layers, followed by max-pooling. The outputs are then flattened and fed through two fully connected linear layers and a final 2-node output layer which determines the predicted label of “airflow” or “touch”. **(C)** Schematic of the SVM classifier consisting of a non-linear embedding using radial basis function (RBF) kernels followed by a linear classifier. **(D)** Sample spectrogram images of “airflow” trials (top) and “touch” trials (bottom) used in the training set of a classifier.

For each labelled baseline period, we compute a spectrogram and use the resulting spectrogram-label pairs to form our dataset. To ensure a large enough dataset for each classifier, baseline spectrograms originating from the same anatomical region in a single session were group together to form one dataset for every session and every nucleus recorded in that session. For each nucleus-session dataset, we trained one instance of a CNN and SVM each (**Figure 3A**). The architecture of the CNN is shown in **Figure 3B** and detailed in Methods Table 1. The first half of the network consists of two successive 2D convolutional layers, followed by a Max Pool layer.

**Table 1.**
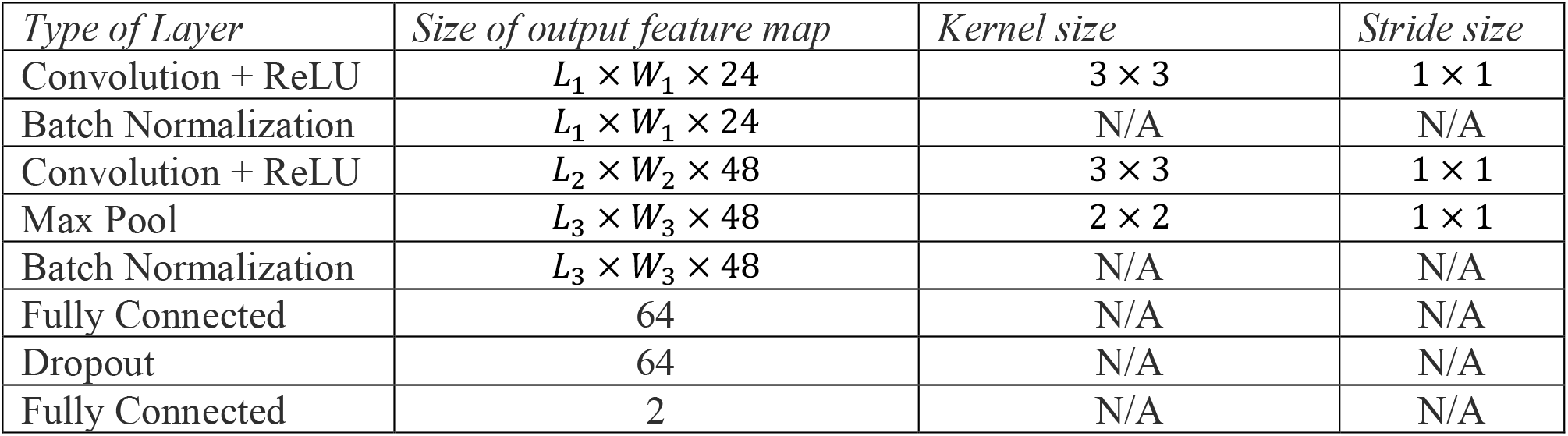
Summary of model architecture. Note that **L**_i_ and **W**_i_ will vary based on frequency bands analyzed and length of input signal.

| Type of Layer | Size of output feature map | Kernel size | Stride size |
| --- | --- | --- | --- |
| Convolution + ReLU | $L_1 \times W_1 \times 24$ | $3 \times 3$ | $1 \times 1$ |
| Batch Normalization | $L_1 \times W_1 \times 24$ | N/A | N/A |
| Convolution + ReLU | $L_2 \times W_2 \times 48$ | $3 \times 3$ | $1 \times 1$ |
| Max Pool | $L_3 \times W_3 \times 48$ | $2 \times 2$ | $1 \times 1$ |
| Batch Normalization | $L_3 \times W_3 \times 48$ | N/A | N/A |
| Fully Connected | 64 | N/A | N/A |
| Dropout | 64 | N/A | N/A |
| Fully Connected | 2 | N/A | N/A |

The convolutional layers learn a set of 48 convolutional kernels (in both time and frequency) which identify distinguishing features of the spectrograms in the training set. These features are then flattened and sent to the second half of the network for classification. The second half of the network consists of two fully connected linear layers which perform linear classification on the features learned from the convolutional layers. In **Figure 3C**, we provide a schematic of the SVM architecture. We train SVM using radial basis functions (RBF) kernel because traditional linear methods failed to discriminate between “airflow” and “touch”, suggesting that a nonlinear embedding is necessary in our context. These modern machine learning methods are useful for detecting patterns in the training set that are not apparent to the naked eye when looking at example trial spectrograms (**Figure 3D**).

### 4. Context can be reliably decoded from baseline LFP in amygdala

Results from training distinct classifiers for each recording session and nuclei are shown in **Figure 4**. For each classifier, we report a summary of the accuracy distribution for a single network computed over 50 instances (more details of the training process are explained in the Methods section). The accuracy for a single instance is calculated as the fraction of correctly labelled spectrograms from the test set. This process is repeated 50 times to generate a distribution of 50 accuracy values for a single classifier. We report the 5^th^, 50^th^ and 95^th^ quantile accuracy for each classifier. There are two classifiers (one CNN and one SVM) for each recording session and each nucleus. Accuracy results using CNNs for all recording sessions across all 3 subjects are shown in **Figure 4A**. Similarly, accuracy results using SVMs for all recording sessions across all 3 subjects are shown in **Figure 3B**.

**Figure 4.**
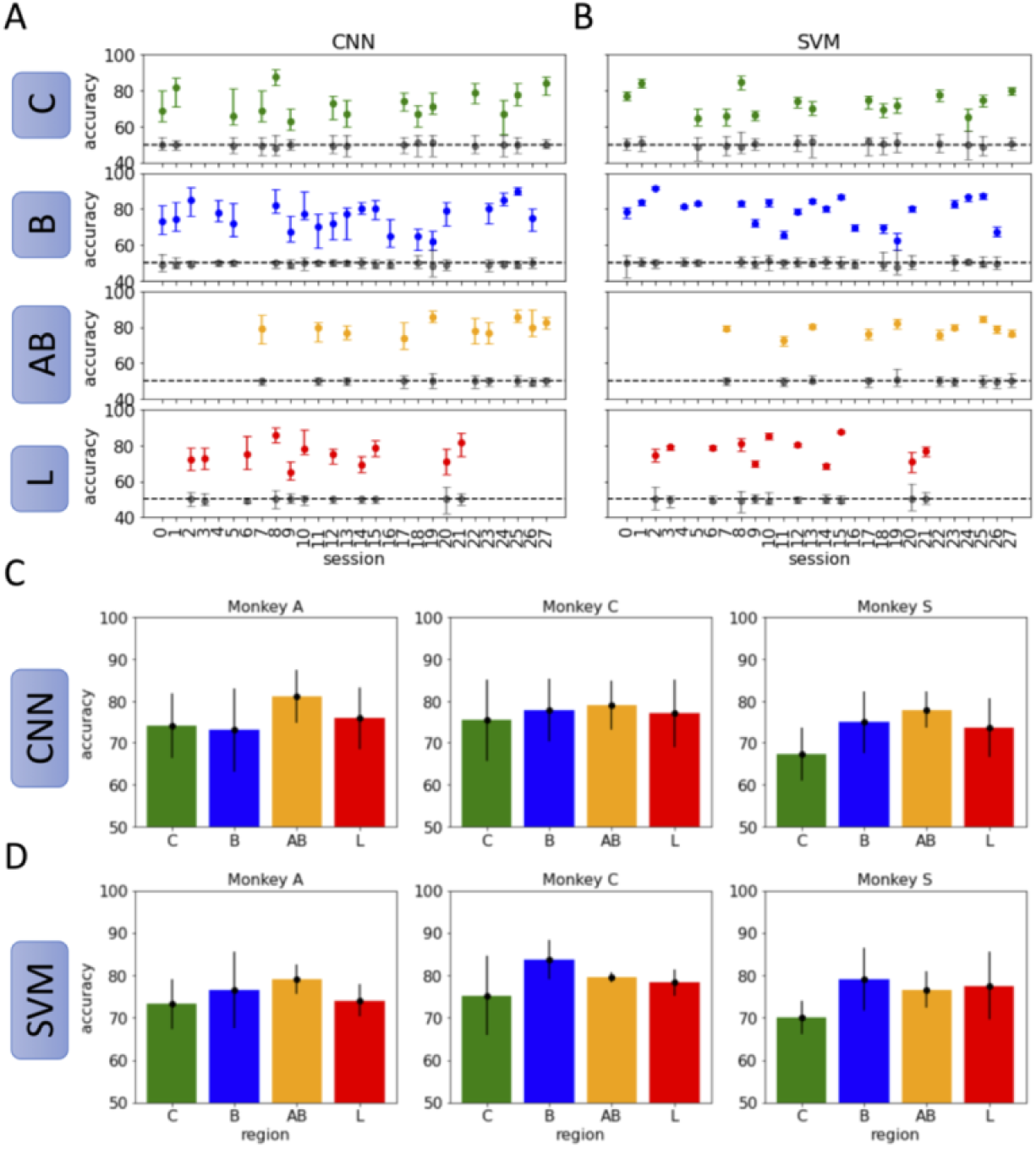
Decoding context reliably from baseline LFP spectrograms. Average accuracy was computed over 50 sample CNNs and SVMs. Trials were randomly reassigned to the training, validation, and testing sets for each sample classifier using an 80-10-10 split. **(A)** Accuracy results for all recording sessions using the CNN classifier. The 50% quantile of accuracy is represented by a dot, with vertical bars reporting the 10% and 90% quantiles. Colors indicate the nucleus in which the recording contacts were located. Gray bars indicate the null distribution obtained from bootstrapping. **(B)** Accuracy results for all recording sessions using an SVM with RBF kernel. Average values and quantiles as in A. **(C)** CNN classification accuracy for each nucleus, averaged over all sessions for three subjects. Gray bars indicate the null distribution obtained from bootstrapping. **(D)** SVM classification accuracy for each nucleus, averaged over all sessions for the same subjects.

Both CNNs and SVM decode context from baseline spectrograms reliably. The distribution of accuracies for each session are shown to be consistently above binary chance (50%). Moreover, to ensure that correlated noise in the datasets was not contributing significantly to network accuracy, we sample from a pseudo-null distribution for each classifier using a bootstrap method. The null distribution is calculated by first shuffling the labels of the training set so that spectrograms are randomly assigned to the “airflow” and “grooming” task equally. The training process is repeated as usual. Using this procedure, we obtain a “null distribution” of accuracies arising from the model performance on true-labelled test set. This gives an estimation for the likelihood of obtaining accuracies better than 50% in a given dataset due to correlated noise. For each recording session and across nuclei, there is no overlap with the null distribution for either classifier. This gives confidence that the classifiers are not decoding context due to chance. The variability in performance across repeated training for each nucleus can be accounted for, in part, by the number of recording contacts present in each nucleus during a recording session. As expected, the accuracy of both CNN and SVM classifiers is positively correlated on the number of contacts present in that region during a recording session (see SI).

### 5. Discriminatory power for context is not nucleus-specific

Next, we determined whether contextual encoding was localized to a particular nucleus within the amygdala. We hypothesized that activity recorded from the basal and accessary basal nuclei would be more reliable for decoding context-related information given that they receive more direct inputs from the prefrontal cortex. Contrary to our expectations we found no difference in decoding accuracy across the nuclei with either classifier type (**Figure 4C, D**). This is confirmed by applying the Kruskal-Wallis H-test (or one-way ANOVA on ranks) across the four conditions. We fail to reject the null hypothesis with p > 0.1 in all three subjects. These results suggest that the context-related signals in the spectrograms of baseline activity do not depend on the hypothesized inputs from the prefrontal cortex, rather, contextual information encoded in baseline arises from inputs that are distributed quasi-equally across the nuclei of the amygdala.

### 6. Discriminatory power relies on information across multiple frequency bands

We next examined the possibility that context is encoded in specific frequency bands. We computed the spike-triggered average (STA) for each nucleus and average over all sessions (mSTA). To identify any frequency-band specific features that might be useful for decoding, we selected the monkey whose recording’s power spectrum showed the largest difference between baseline activity of airflow and touch blocks. The differences in mSTA between puff and touch trials for Monkey A are show in **Figure 5A** (left). The average conditional expectation of the STA power spectrum given that a spike occurred during baseline is also shown in **Figure 5A** (right). This conditional expectation is calculated by computing the power spectrum of the LFP signal around each baseline spike and taking the average across all spectra. From this analysis we see two prominent differences in power between airflow and touch baseline trials, 10-17Hz and 17-25Hz. The lower band having more power during the baseline period between the “touch” trials across multiple (but not all) session and the 17-25Hz band having more power during the baseline period between “airflow” trials consistently across recording sessions. With these bands identified, we re-trained 50 instances of each network using only time-frequency data from the lower band (10-17Hz), the higher band (17-25Hz) or both (10-25Hz). While the average spectrum suggests these bands are important, accuracy of the network on the full spectrogram out-preforms those networks trained on these bands alone (**Figure 5C**). Power spectra were not the same across all three subjects. However, given that in the animal where the effect was strongest there was no benefit of selecting a particular frequency band, we conclude that contextual encoding is not restricted to a particular frequency band.

**Figure 5.**
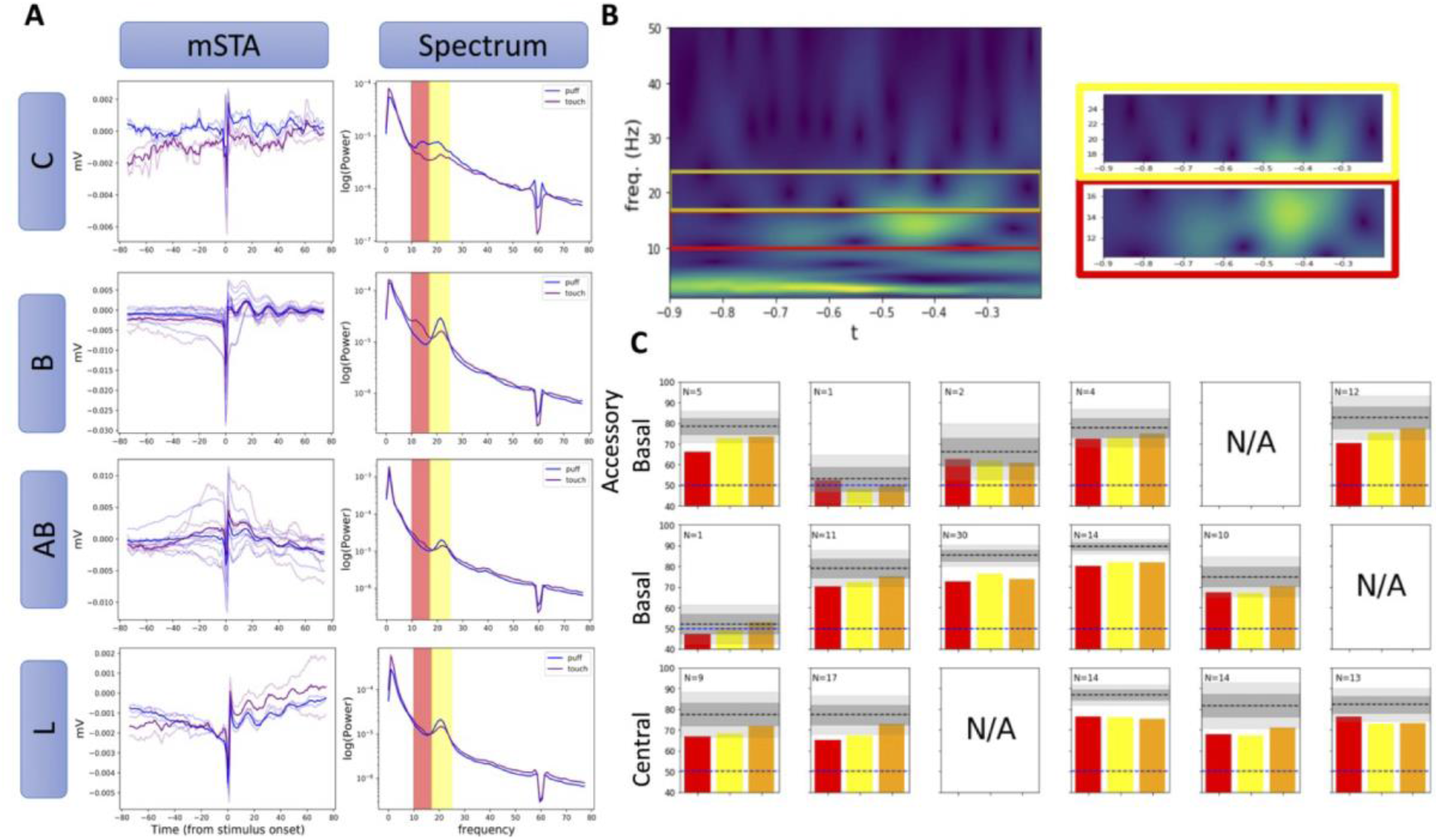
Discriminatory power is distributed across a wide range of frequency bands. **(A)** Left: Mean Spike-Triggered Average (mSTA) traces for “airflow” (blue) and “touch” (purple) computed for ±80 ms relative to spikes occurring during baseline. mSTA is computed by averaging STAs over all cells in the same nucleus. The number of visible lines corresponds to the stable cells in each nucleus used to compute mSTA. Pale lines are single STA traces and dark lines are the mean STA. Right: Comparison of average power spectra of STA traces for airflow and grooming blocks. The two spectra show differences in the 10-17 Hz (red) and 17-25 Hz (yellow) bars. **(B)** A trial spectrogram illustrating the power in the 10-17Hz (red) and 17-25Hz (yellow) frequency bands. **(C)** Accuracy of CNNs trained on spectrograms restricted to only 10-17 Hz (red), 17-25Hz (yellow) and 10-25 Hz (orange) bands. Plots are organized by nucleus (rows) and recording session (columns), with number of recording contacts in each nucleus displayed in the top left corner. The horizontal black dotted line represents the mean accuracy of the network trained on the full spectrogram. The horizontal blue line represents theoretical chance at 50%. The gray bars correspond to 1 and 2 standard deviations about the mean.

## DISCUSSION

In neurophysiology, “baseline” refers to the spontaneous ongoing activity of neurons in the absence of the organism’s engagement with external stimuli or task variables. External stimuli and cognitive processes shift the brain away from baseline toward task-specific or stimulus-specific functional states. However, at the cessation of the external stimulus or the completion of the cognitive process, the brain is expected to return to the same baseline. Here we show that this is not always the case; when the context is different the baselines are also different. Although few studies addressed directly the content of the baseline activity, the discovery of intrinsic dynamics of the brain, such as memory replay (Skaggs and McNaughton, 1996), the default mode network (e.g., Raichle, 2015), and off-task cognitive process (Kucyi et a., 2023), raised the possibility that the baseline activity carries relevant and decodable information (Kaefer et al., 2022).

As context is often signaled to the primate amygdala by the prefrontal cortex (Rigotti et al., 2010; Saez et al., 2015), it is expected that the nuclei that receive direct prefrontal inputs would show the strongest context-related activity. Specifically, the basal and accessory basal nuclei of the amygdala receive monosynaptic inputs from multiple prefrontal areas whereas the lateral and the central nuclei are connected to the prefrontal cortex through multi-synaptic pathways (Ghashghaei et al., 2007; Barbas et al., 2011; Cho et al., 2013; Romanski, 2007). Contrary to this prediction, both CNN and SVM decoded with similar accuracy context-related signals from the LFPs in all the nuclei of the amygdala. It is unlikely, therefore, that the spectrograms of baseline activity are shaped by direct synaptic currents transmitted from prefrontal areas to select nuclei of the amygdala. It is more likely that a broader, global phenomenon, such as the autonomic state of the animal is signaling context to the amygdala. Indeed, we report significantly different autonomic states during the airflow and the touch blocks (**Figure 2**). During the airflow blocks the monkeys are by themselves in a booth and while they receive innocuous airflow stimuli directed at their face, they are alert, attentive, and highly responsive to external stimuli. During the touch blocks, when a bonded and trusted human grooms their face the monkeys close their eyes and relax their facial musculature. The groomer’s presence is marked by reductions in heart rate and increases in heart rate variability that indicate lower sympathetic tone and higher parasympathetic tone (Berntson et al., 1993; Quigley and Barrett, 2014). The autonomic state of the body is transmitted to multiple areas of the brain through interoceptive pathways (Craig, 2002; Craig, 2003; Berntson and Khalsa, 2021). It is possible, therefore, that the baseline activity across all nuclei of the amygdala is shaped by interoceptive inputs that were widely different during the airflow and touch blocks. Similar modulation of the baseline activity by interceptive inputs have been documented in mice (Livneh and Andermann, 2021). Whether LFPs in different nuclei of the amygdala carry different or similar signals, it remains unclear how context-related or interoception-related features of LFPs might contribute to decoding accuracy.

Our data show that all frequencies contribute to decoding context from the baseline LFPs. Indeed, there are no prior studies in the primate amygdala that would bias our expectations for any of the frequency bands. In the cortex, bottom-up and top-down signals travel through brainwaves of different frequencies (Buschman and Miller 2007). For example, executive functions, localized typically to the prefrontal cortex of primates, require an interplay between the high frequency, gamma (30–100 Hz) oscillations localized to the upper cortical layers (layers 2 and 3) and low-frequency oscillations in the alpha and beta bands (10–30 Hz) localized to deep cortical layers (layers 5 and 6) (Miller et al., 2018). The theta rhythm (8-12 Hz) so prevalent and so broadly explored in rodents appears to be less tractable in non-human primates (Abbaspoor et al., 2023). These low-frequency bands have been observed in primates in response to the predictable stimuli. The power in the high-frequency bands is enhanced in response to unpredictable stimuli (Bastos et al., 2020). All oscillations, however, can change the functional connectivity between brain areas that support different cognitive components of complex tasks (Pinotsis et al., 2019). We show that access to broadband frequency information across the entire baseline trial improves decoding accuracy significantly. These finding suggest that the dynamics of context-specific baseline activity are not characterized by sustained localization of activity in one frequency band or another. It is more likely that these context-dependent states give rise to more complex, transient dynamics involving the interplay of multiple frequency bands. One possibility is that the network dynamics emerging from context-dependent states include short “bursts” of synchronous activity that may bifurcate to multiple frequency bands and dissipate.

Indeed, baseline spectrograms show evidence of complex spatiotemporal patterns with transient bursts (localized bright spots), broad band power (vertical swaths) and apparent bifurcations (holes) (**Figure 3D**). Such higher-order, transient dynamics would require knowledge of broadband frequency information across time to accurately decode. Biologically plausible mechanisms that could give rise to such patterns remain obscure. Further analysis of these spectrogram features and how they contribute to decoding accuracy is ongoing.

### Comparison with previous work

A limitation of the study was the lower decoding accuracy compared to single unit population activity during the same baseline periods (Martin et al., 2023). This may be related to the short baseline periods (approximately 500 ms) and the low number of trials (about 400 trials for training) available for session and subject specific analysis. In the single unit approach, stable cells identified across all recording sessions and subjects (for a total 237 units) were used to train a single SVM. These cells were randomly sampled with replacement to train the SVM and it was determined that a population of 127 single units were predictive of context at the 95% confidence level. This population size of single units is far greater than the number of stable cells recorded in an individual session and makes no distinction between nuclei. Given that our LFP approach is nucleus-, session-, and even subject-specific, it is to be expected that decoding accuracy with LFP is lower than with single unit activity.

### Comments on ML methodology

In this paper, we have not aimed to train classifiers that are generalizable. Our classifiers are trained on data obtained from different subject and sessions when the linear probes recorded neural activity from different nuclear subdivisions of the amygdala. In fact, the classifiers do *not* generalize when applied to data obtained from other sessions, which was expected based previous work that mapped dissociable functions to different mesoscale subregions of the amygdala (Morrow et al., 2019). For the purpose of this paper, our (non-generalizable) approach was sufficient to conclude that contextual information is present in baseline LFP. In the future, we plan to study other ML architectures that might be more robust to variability across subjects and small changes in probe placement, possibly by incorporating biophysics-informed ML methods. We suspect developing ML architectures that disentangle biological sources of variability from measurement-specific variability will be necessary for generalizability.

It remains to determine which features of LFPs were used by our classifiers to achieve their performance. We expect future work focusing on baseline activity in the amygdala will shed light on how this often-ignored feature of brain activity holds specific information about context, interoception, and other aspects of brain states. If we are able to isolate those features of LFP were used by our ML classifiers to achieve their performance, then our methods – in addition to being effective information detectors – would be useful for generating new hypotheses and may directly contribute to a clearer picture of contextual information coding in the amygdala and elsewhere. A closely related question is why spectrograms are so effective in reflecting contextual information. Indeed, we have applied similar methods to “raw” LFP time series data, and the results show that time-frequency information improves decoding accuracy markedly (SI). These questions are the subject of on-going investigation.

## MATERIALS AND METHODS

### Experiment

For a detailed description of the experiment and data collection, see the Methods section of Martin et. al. (2023).

### Materials

All models and data analysis were run with Python using popular open-source packages like *Pytorch-1*.*11*.*0, numpy-1*.*22*.*4), scipy-1*.*12*.*0, and scikit-learn-1*.*1*.*1*. We have released all relevant source code at https://github.com/anaucoin/Aucoin-ML-LFP-2024. Data available upon request.

### Heartrate and Respiratory Sinus Arrhythmia

Instantaneous heartrate values were computed from the inverse of the duration between two heartbeat times (IBI). Values above 240 BPM or below 40 BPM (IBI below 250ms and above 1500ms) were removed. All noise and movement artifacts identified were also removed. Heartbeat values were interpolated to a 1ms timescale using a modified Akima cubic Hermite polynomial.

Heartrate variability (Berntson et al., 1993; Sztajzel 2004) was calculated using a spectral density estimation method. Spectral power density was computed from the cleaned heartbeat times using a multitaper method in sliding windows of 60 seconds and an overlap of 3s. A total of 7 Slepian tapers were used for smoothing. The spectra in each time window were normalized to have unit area between 0.25 Hz and 0.5 Hz, corresponding to respiratory rates of 15 and 30 breaths per minute. We define respiratory strength in each time window as the average power at the peak ±halfwidth for peaks occurring between 0.25 Hz and 0.5 Hz. In time windows with no peaks, the average power across the entire 0.25 to 0.5 Hz window was used. RSA strength was then normalized to the median power across all time windows

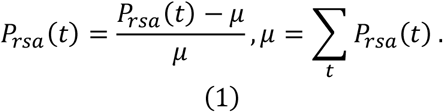

### Spectrogram computation

To compute trial spectrograms, we use a Continuous Wavelet Transform (CWT) method. Unlike traditional spectral methods like Fast-Fourier Transform (FFT), which collapse time, CWT provides a trade-off between spectral and temporal resolution. CWT is a natural choice as we do not expect trial LFPs to be stationary. For a given signal *x*(*t*), the CWT 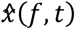 is defined as 

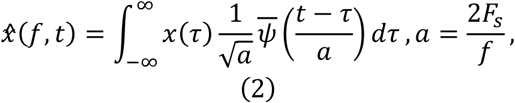

where 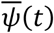 denotes the complex conjugate of the wavelet function *ψ*(*t*), and *F*_*s*_ is the sampling rate of the given input signal *x*(*t*).

For each trial baseline LFP, we computed the CWT using a Complex Morlet Wavelet

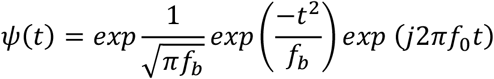

where *f*_0_ is the central frequency and *f*_*b*_ is the bandwidth. All spectrogram images were computed using the *scipy*.*signal*.*cwt* function with ‘Morlet2’ wavelet and a standard central frequency of 5 for a good balance of temporal and frequency resolution. To reduce model training computation time, we limited the input image size by considering only frequencies in the range of 1-50 Hz for each spectrogram. Similar networks were trained with frequencies up to 100Hz and showed no significant increase to model accuracy.

### Convolutional Neural Network

Convolutional Neural Networks (CNNs) are deep multi-layer networks widely used in computer vision and image classification tasks. The simple CNN architecture used in our paper can be thought of in two parts: (1) a convolution operator and (2) a linear classifier. The layers in the convolutional part of a CNN typically include three operations:

1. Convolution
2. Non-linear activation
3. Pooling

though not all operations need be in every “layer” of the network. The convolutional layer involves convolving an input image *x* with a collection of K kernels of size *s* × *s. s* is typically chosen to be small to preserve locality. In our network, *s*=3. The kernels are learned by the network during training and output a collection of K local features. These features are then passed through a non-linear activation function (typically ReLU) and then through a pooling operation (such as Max or Average Pooling). The non-linear activation can be thought of as thresholding and increasing the receptive field in a biological sense, and the pooling operation serves to further reduce dimensionality. These steps can be repeated to increase the depth and complexity of the network. The result of these successive operations is a collection of features extracted from each input image. These features are then flattened into a single vector and used as input into the linear classifier part of the CNN. This second half of the network consists of a series of fully connected linear layers, which compose a linear map that transforms the flattened features into an output vector *y* of length *C*_*n*_, the number of classes. Each 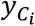 describes the probability that the input image *x* belongs to class *C*_*i*_. The weights and biases of the linear map are optimized during training.

The exact model architecture and parameters used in this analysis, can be found in Table 1. For each recording session, and for each nucleus recorded in that session, we trained a CNN to classify spectrograms of the baseline LFP signals as “airflow” or “touch”. The baseline spectrograms for all channels in a particular anatomical region were used as inputs into the network for the training, validation, and testing stages. Bootstrapping (random sampling with replacement) was used to sample the “touch” images so that there was an equal number of “touch” and “airflow” images in the dataset. The data was then split into training, validation, and testing subsets with an 80-10-10 split. An equal class representation was ensured in each split. Each input image was normalized using the “MinMaxScaler” from *scikitlearn* package fit only on the training set to prevent erroneously giving the network information about the entire dataset. The network parameters were updated through backpropagation using the Adam optimizer and a cross-entropy loss function was used.

The data was fed into the model in mini batches of size 20 and trained for 40 epochs. This means that before the model saw 20 spectrogram images before updating the model parameters and saw all spectrogram images a total of 40 times. During training, the model was shown spectrogram images from the validation set to track over-fitting of the model. If the accuracy of the model improved on both the training set and validation set, the model parameters were saved as the current best instance of the model. Once all 40 epochs were complete, the best model (the model with the lowest validation accuracy), was used in the testing stage of the model. To analyze the accuracy and robustness of each model, we repeated this procedure 50 times. We reinitialized the model, randomly assigned spectrograms to the training, testing, and validation sets, trained the new model and recorded the accuracy on the test set. The accuracy reported for each model is the average percentage of correctly classified images over 50 model initializations. The robustness is tracked by the 5^th^ and 95^th^ quantiles.

### Support Vector Machine

Support Vector Machines (SVM) are a popular supervised learning method for classification and regression. They have been shown to be effective even in high dimensional settings and settings where the number of dimensions is much higher than the number of samples. SVMs construct a high-dimensional hyperplane, called the decision boundary, to separate training data by their labeled class. This hyperplane is chosen to maximize the margin, or distance between the decision boundary and nearest data points. In the linear classifier case, this is equivalent to solving the following optimization problem:

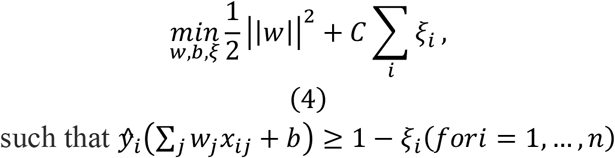

In the case where a linear decision boundary is insufficient, the linear classifier can be made nonlinear through a non-linear kernel function. The kernel functions embed the data *x*_*i*_ into a new vector space called the feature space. This procedure is sometimes called the “kernel trick”. The new optimization problem replaces *ŷ*_*i*_(Σ_*j*_ *w*_*j*_*x*_*ij*_ + *b*)in (4) with **ŷ**_*i*_, where *ψ* is the choice of kernel function. In practice, the dual formulation of this optimization problem is used to avoid explicitly mapping the data points into the feature space. This helps to better scale the computational efficiency and memory usage, especially when the number of features in the data is high.

To train the SVM, we used svm.LinearSVC and svm.SVC from the popular *scikitlearn* package. We trained a non-linear classifier with radial basis kernel functions using the dual form for efficiency. The input data into the SVM are the flattened spectrograms labelled as “airflow” or “touch”. The dimension of the input data is *T* × *F* where T is the number of time points and F is the length of the frequency space discretization used. F and T will vary on the frequency bands of interest and the length of LFP baseline for that session. Training, validation and testing of the SVMs were the same as in training of CNN.

### Spike Triggered Average

In each session and each recording electrode, spike-triggered averages (STA) were computed for each by selecting a window of ±80ms around those spikes occurring during the baseline period and taking the average over all windows. Power spectrum of the STA were computed using Welch’s method with Hanning tapers, a segment length of 1ms and 50% overlap. Mean STA (mSTA) and average STA spectrums were taken as the average STA and average spectrum over all electrodes in the same nucleus.

## ACKNOWLEDGEMENTS

AA was supported in part by NSF grant DMS-1937229. KL was supported in part by NSF grant DMS-1821286. KMG was supported by R01MH121009. We thank Dr. Anne Martin, Michael Cardenas, Rose Andersen, Archer Bowman, and Elizabeth Hillier for data collection, data organization, the analysis of single unit and heart rate data, and the initial processing of the LFPs. Michael Cardenas helped with the RSA calculation. Dr. Francesco Battaglia offered useful suggestions for improving the early versions of the manuscript.

## SUPPORTING INFORMATION

### Baseline selection criteria

Baseline LFP was selected from a stable time window of the interstimulus interval (ISI) between two stimuli of the same type. The ISI was defined as the period occurring 200ms after stimulus offset and 200ms before stimulus onset. For each recording session, all ISI signals were trial averaged and the standard deviation for each timepoint was calculated. Baseline LFP for each trial was chosen by inspecting the trial-averaged ISI and determining a time window with low trial-wise variability.

**Figure S1.**
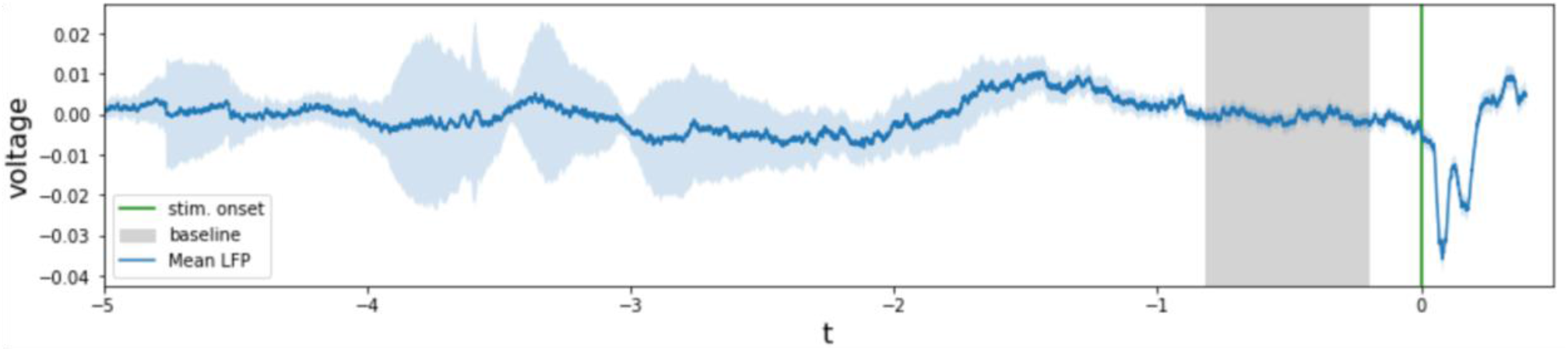
Selecting stable baseline trials. An example of the baseline LFP selection criteria for a single recording session. The solid blue line is the trial-average of the signals during each ISI. The vertical green line is stimuli onset. The light blue shading is 2 st. dev. of the mean. The gray box is the stable time window chosen as the baseline.

### Linear SVM fails to discriminate context reliably

**Figure S2.**
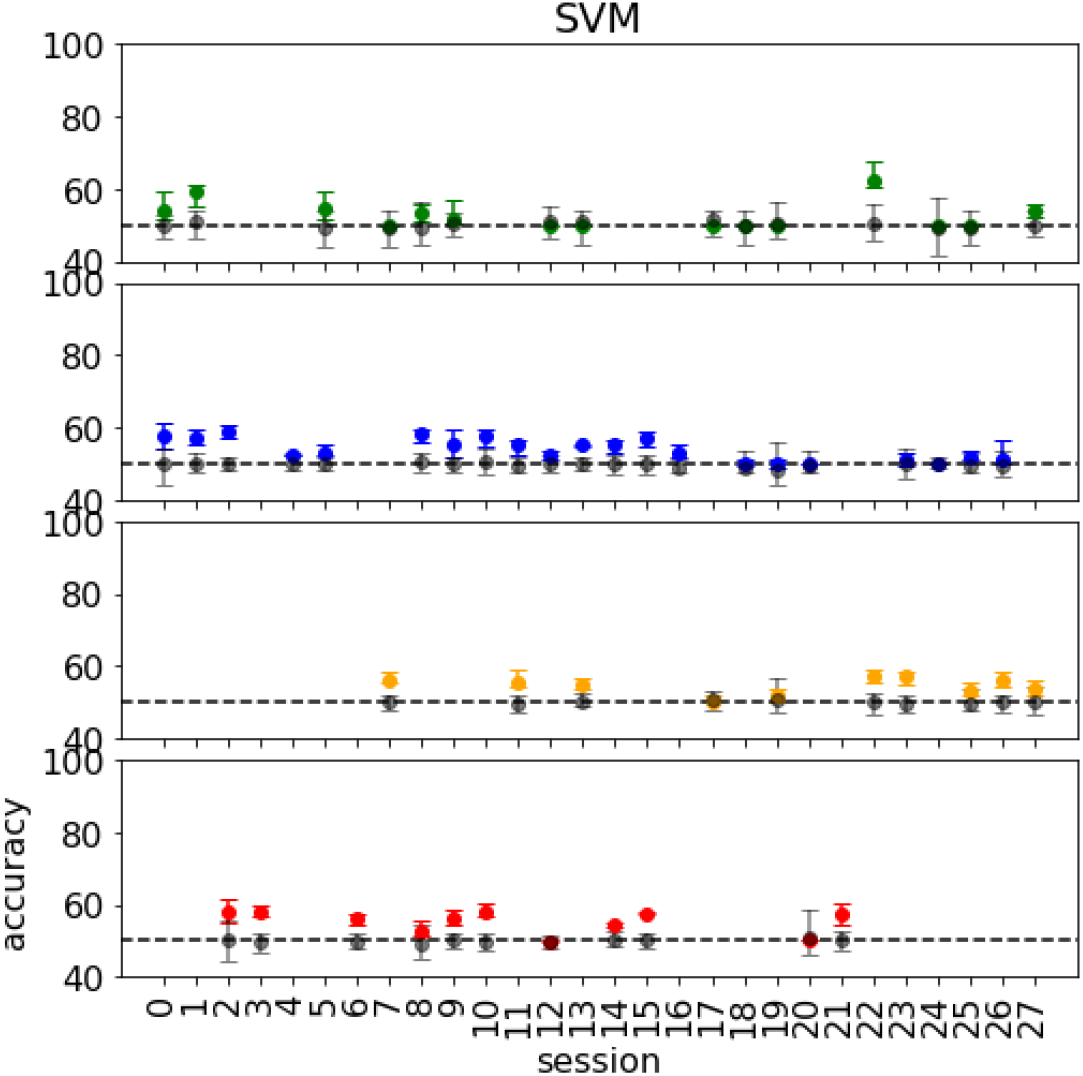
Decoding context fails with linear SVM. Average accuracy was computed over 50 sample linear SVMs. Trials were randomly reassigned to the training, validation, and testing sets for each sample classifier using an 80-10-10 split. Accuracy results for all recording sessions using the CNN classifier. The 50% quantile of accuracy is represented by a dot, with vertical bars reporting the 10% and 90% quantiles. Colors indicate the nucleus in which the recording contacts were located. Gray bars indicate the null distribution obtained from bootstrapping.

Spectrogram classification failed on SVM without using the kernel trick. This is likely because the time-frequency features are highly correlated and require a non-linear mapping to improve separability. Results from the linear SVM classification are shown in **Figure S2**. Average accuracy across nuclei and sessions rarely exceeds 60% and often is not statistically different from the null distribution. Overall, the linear SVM classification is less reliable than performance from both SVM with kernel trick and the CNN, both of which leverage non-linear transformations of the data before performing classification, indicating that the non-linear embedding is necessary for classification.

### Classification accuracy depends on dataset size

Variability across recording sessions and nuclei is due, in part, to the number of recording electrodes present during recording sessions, which determines the total amount of data. Machine learning methods are notoriously data-hungry, and typically require a large amount of data to adequately learn a particular task. **Figure S3** show the classification accuracy as a function of the number of recording electrodes in a particular nucleus during a recording session. Results for all three subjects are shown. For both CNN and SVM, the accuracy of days with a single recording electrode is below 60% but quickly improves with the addition of more data. Both classifiers also show performance generally plateauing between 70-85% which can be achieved with 5 or more contacts present. This exploration explains why performance of the classifier on the central nucleus is (slightly) lower than other nuclei across subject. The relative size of the central amygdala compared to other nucleus is much smaller, limiting the total amount of electrodes that can be present in central amygdala in any given session. Nevertheless, we still see both SVM and CNN classifiers performing better than chance when data from more than one recording electrode is available.

**Figure S3.**
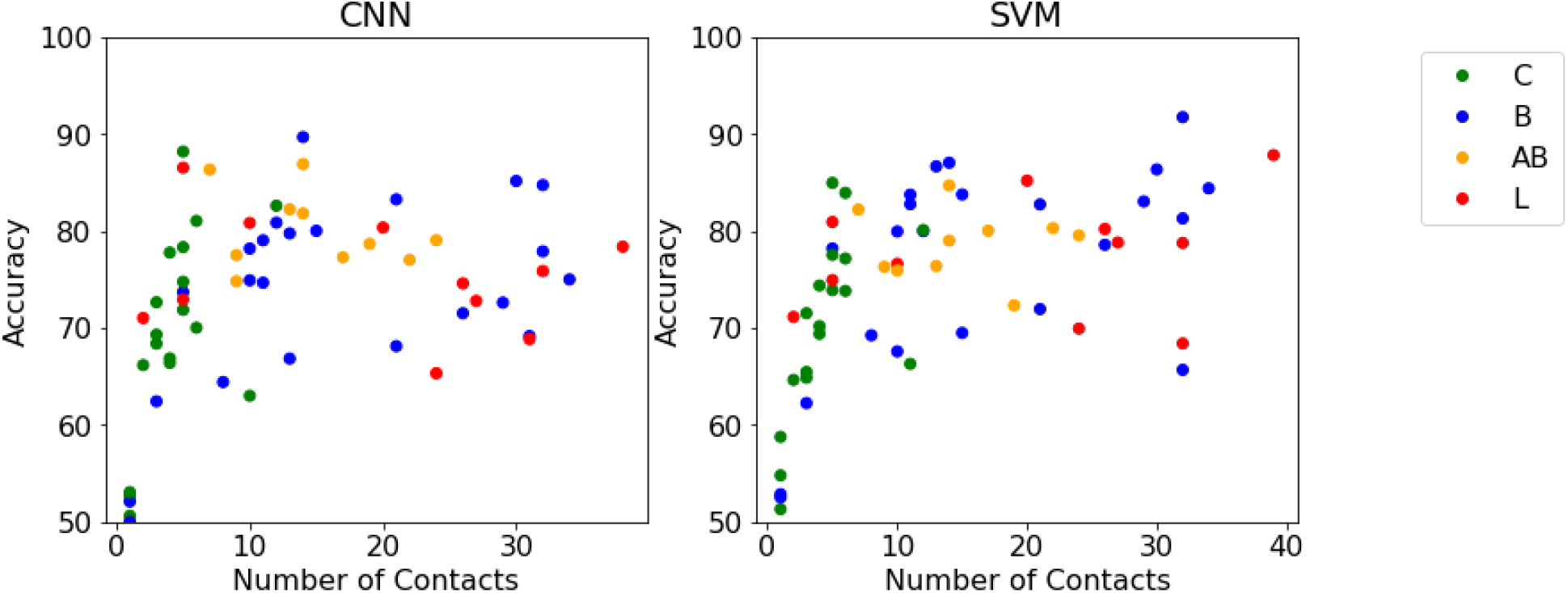
Decoding accuracy is a function of data availability (number of recording electrodes). Classification accuracy for one sample of CNN (left) and SVM (right). Each dot represents the accuracy of a single classifier trained on data from one session and nucleus.

### Computational cost of CNN and SVM

Accuracy results in the main body of the text suggest that although SVM and CNN perform the same on average, the SVM classification exhibits less variability across the 50 realizations (evidenced by the narrower confidence intervals). Readers may be tempted to assume that SVM implementation should therefore be preferred over CNN. However, the choice in classifier may be better suited by the amount of data available. **Figure S4** shows the computational cost of implementing both CNN and SVM as a function of the number of recording electrodes. As mentioned above, the number of recording electrodes will strongly influence the amount of data in the training set. For small amount of data (<4 electrodes), SVM is the more efficient implementation. However, overall, SVM scale quadratically with the number of electrodes (as expected) while the CNN scales linearly. For moderate and large amounts of data, the CNN is more efficient.

**Figure S4.**
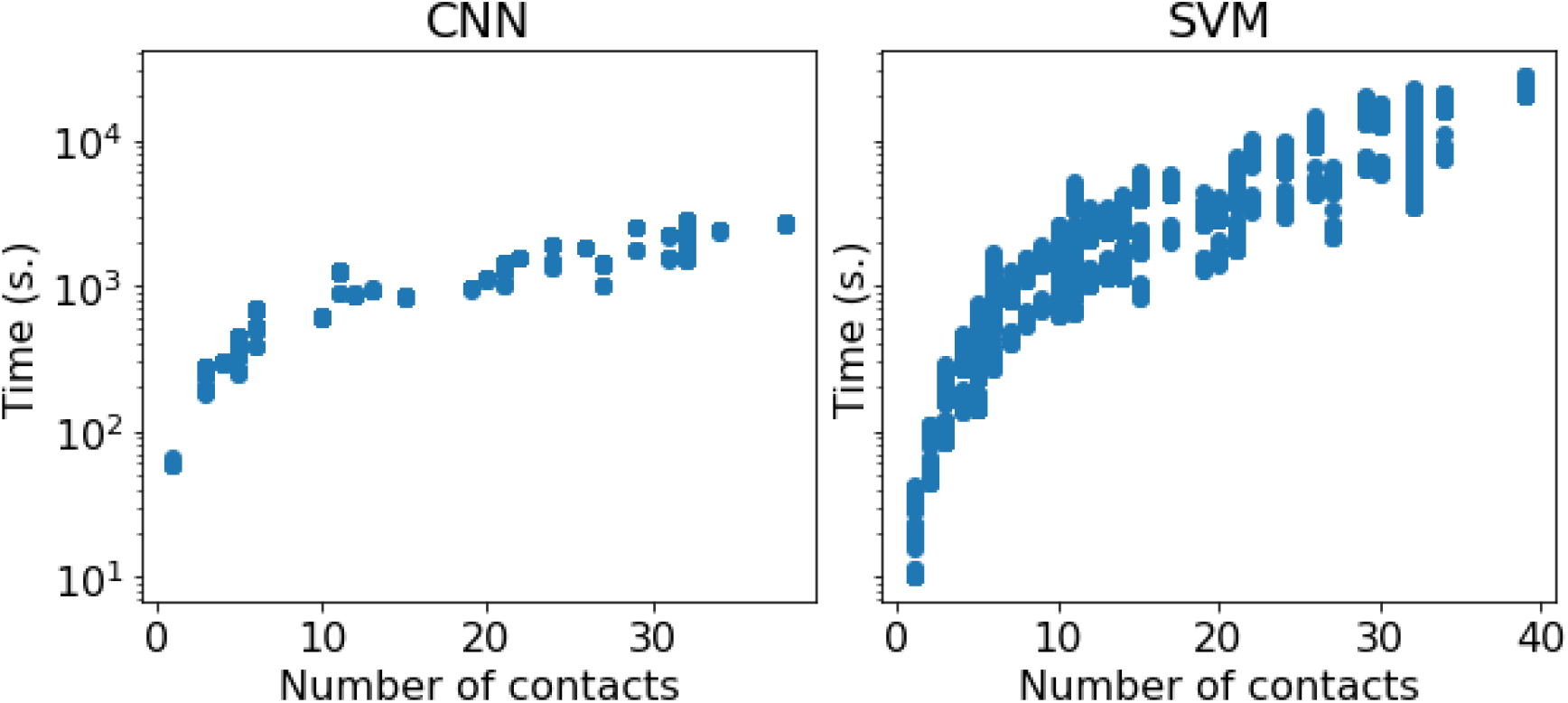
Computational costs of CNN and SVM. The computational time for training a single classifier for CNN (left) and SVM (right) as a function of the number of recording electrodes (which most influences training dataset size). Each dot represents a single realization of the classifier.

### Classification performance decreases dramatically using raw time-series alone

The methodology presented in the main text uses trial spectrograms as the feature space for decoding. As a first step in our analysis, we tried traditional classification methods, like SVM, using the raw time series data. Though these methods were able to discriminate between airflow and grooming contexts, the accuracy results were little better than chance. **Figure S5** shows the accuracy of SVM decoding using the raw time series trials compared to spectrograms. Decoding accuracy noticeably improves across all subjects and nuclei when using time-frequency spectrograms (Monkey A: 10% increase, Monkey C: 18%, Monkey S: 20%)

**Figure S5.**
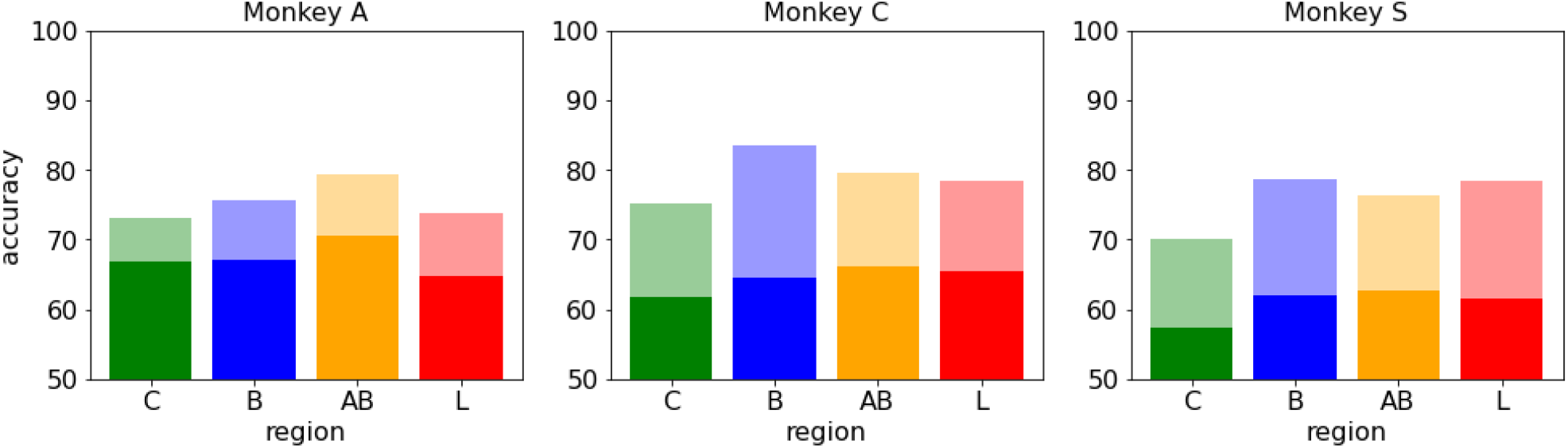
Comparison of SVM using raw timeseries data over spectrogram. SVM classification accuracy for each nucleus, averaged over all sessions for the same subjects (Monkey A (left), Monkey C (middle), Monkey S (right)). More saturated colors indicate the SVM accuracy using raw time series trials. SVM decoding accuracy using trial spectrograms (reported in the main text) is shown in more transparent color for comparison.

